# An N-terminal Fusion Allele to Study Melanin Concentrating Hormone Receptor 1

**DOI:** 10.1101/2021.04.23.440964

**Authors:** Kalene R. Jasso, Tisianna K. Kamba, Arthur D. Zimmerman, Ruchi Bansal, Staci E. Engle, Thomas Everett, Chang-Hung Wu, Heather Kulaga, Randal R. Reed, Nicolas F. Berbari, Jeremy C. McIntyre

## Abstract

Cilia on neurons play critical roles in both the development and function of the central nervous system (CNS). While it remains challenging to elucidate the precise roles for neuronal cilia, it is clear that a subset of G-protein-coupled receptors (GPCRs) preferentially localize to the cilia membrane. Further, ciliary GPCR signaling has been implicated in regulating a variety of behaviors. Melanin concentrating hormone receptor 1 (MCHR1), is a GPCR expressed centrally in rodents known to be enriched in cilia. Here we have used MCHR1 as a model ciliary GPCR to develop a strategy to fluorescently tag receptors expressed from the endogenous locus *in vivo.* Using CRISPR/Cas9, we inserted the coding sequence of the fluorescent protein mCherry into the N-terminus of *Mchr1*. Analysis of the fusion protein (^mCherry^MCHR1) revealed its localization to neuronal cilia in the CNS, across multiple developmental time points and in various regions of the adult brain. Our approach simultaneously produced fortuitous in/dels altering the *Mchr1* start codon resulting in a new MCHR1 knockout line. Functional studies using electrophysiology show a significant alteration of synaptic strength in MCHR1 knockout mice. A reduction in strength is also detected in mice homozygous for the mCherry insertion, suggesting that while the strategy is useful for monitoring the receptor, activity could be altered. However, both lines should aid in studies of MCHR1 function and contribute to our understanding of MCHR1 signaling in the brain. Additionally, this approach could be expanded to aid in the study of other ciliary GPCRs.

## Introduction

G protein-coupled receptors, GPCRs, comprise a large family of signaling proteins that function throughout the body (Drake, Shenoy, & Lefkowitz, 2006; Marinissen & Gutkind, 2001). Many of these receptors are still considered “orphan” or lack identified ligands, and the cellular localization for many is also unknown due to a lack of good antibodies or epitope availability within *in vivo* contexts (Michel, Wieland, & Tsujimoto, 2009). Of the non-olfactory GPCRs, approximately 10% have been found to preferentially localize to a signaling compartment called the primary cilium (Berbari, Johnson, Lewis, Askwith, & Mykytyn, 2008; Kroeze, Sheffler, & Roth, 2003; McIntyre, Hege, & Berbari, 2016; Schou, Pedersen, & Christensen, 2015). These small microtubule-based appendages serve as signaling centers on most mammalian cell types, including neurons throughout the central nervous system (Berbari, O’Connor, Haycraft, & Yoder, 2009). Approaches and tools are needed to reveal the significance of GPCR sub-cellular localization and subsequent impacts on *in vivo* signaling and physiological roles.

Mouse alleles that express fluorescently tagged ciliary GPCRs have the potential to be a useful alternative to antibody labeling. However, fluorescently tagged ciliary GPCR mouse models are limited. It is challenging to produce fusion alleles that do not interfere with ligand binding, cilia localization, or downstream signaling. Intracellular C-terminal tags could interfere with intracellular signaling proteins or with ciliary localization sequences in the third intracellular loop (Berbari, Johnson, et al., 2008). Cilia^GFP^ mice express the GPCR, somatostatin receptor 3 (SSTR3) tagged with GFP on the C-terminal. While SSTR3-GFP cilia localization is retained, male mice are infertile suggesting its expression impacts cilia or flagella function (O’Connor et al., 2013). The GFP tag in another ciliary GPCR reporter, rhodopsin-GFP, is also located on the C-terminal and has normal expression patterns in mice but homozygosity results in retinal degeneration (Chan, Bradley, Wensel, & Wilson, 2004). In addition to tagged GPCRs, transgenic cilia membrane associated reporter alleles like Arl13b-mCherry or Arl13bmCherry-GECO1.2 are broadly expressed perhaps outside of their endogenous patterns and levels, which may also influence cilia associated phenotypes and signaling processes (Bangs, Schrode, Hadjantonakis, & Anderson, 2015; Delling et al., 2016). Thus, mice with fusion tagged GPCRs whose expression is driven from the endogenous locus may serve as useful tools to understand the role that ciliary localization plays in receptor signaling. However, they should be carefully screened for both proper expression and function. These types of tools could provide several additional advantages over antibody labeling. One benefit of fusion alleles is the ability to assess the subcellular localization of GPCRs when antibodies are not available or under any conditions that may inhibit antibody binding, such as ligand binding or epitope altering post-translational modifications associated with signaling (Patwardhan, Cheng, & Trejo, 2021). Extracellular fluorescent fusion proteins are also beneficial for monitoring dynamic localization in live imaging approaches and monitoring internalization (Jenkins et al., 2011; McEwen et al., 2007).

We chose to generate an N-terminal fusion GPCR allele using CRISPR/Cas9 of the known ciliary melanin concentration receptor 1 (MCHR1) and directly assess the impact of this fusion allele on receptor expression, localization and function. While MCHR1 is enriched in primary cilia, many questions remain about how MCHR1 function is impacted by its ciliary localization. In rodents, MCHR1 is the only known receptor for the neuropeptide melanin-concentrating hormone (MCH). MCH is produced by neurons found in both the hypothalamic zona incerta and the lateral hypothalamus that project widely throughout the brain (Bittencourt et al., 1992; Engle et al., 2018; Saito, Cheng, Leslie, & Civelli, 2001). The broad innervation of MCH fibers and expression pattern of MCHR1 allows it to impinge on several different behavioral processes including feeding, drinking, addiction, mating, sleep, and maternal behaviors (Diniz & Bittencourt, 2017). While it is clear that MCH pathway manipulation in various brain regions impacts behavior, how this neuropeptide and its receptor act at the cellular level to influence this wide array of behaviors remains unclear. For example, does MCHR1 dynamically change its cilia localization upon ligand addition? Some ciliary GPCRs and signaling machinery have been found to dynamically localize to the compartment in heterologous expression systems and cell lines, but it is unclear if they do so *in vivo* (Green et al., 2016; Shinde, Nager, & Nachury, 2020; Ye, Nager, & Nachury, 2018). How does ciliary localization impact G protein-coupling and neuronal activity? How does ciliary localization of MCHR1 impact behaviors like feeding, sleep and reward? New tools to study the MCHR1 receptor would help answer these questions and be valuable for a wide variety of research involving not only MCH signaling but may also reveal general themes in regards to neuronal ciliary signaling.

Using a CRISPR/Cas9 mediated approach, we have generated a novel fluorescently N-terminal mCherry tagged *Mchr1* reporter allele that allows for monitoring of MCHR1 localization during development, as well as a new *Mchr1* knockout allele. Here we demonstrate that the resulting fusion protein, ^mCherry^MCHR1, faithfully matches the known neuroanatomical expression pattern and subcellular localization of MCHR1. We further show that an 8bp deletion including the *Mchr1* start codon is sufficient to abolish MCHR1 detection by immunofluorescence and alter synaptic activity. We also provide evidence that in the homozygous condition, expression of ^mCherry^MCHR1 also slightly reduces synaptic activity, revealing the need for properly controlled approaches when tagging ciliary proteins. Future studies will determine how certain behaviors may impact the subcellular localization of MCHR1 as well as the signaling ability of ^mCherry^MCHR1.

## Results and Discussion

### Generation of ^mCherry^MCHR1 reporter and MCHR1 knockout

Fluorescent epitope tags provide additional means for the study of GPCRs and for understanding GPCR subcellular localization (Ehrlich et al., 2018; Lu et al., 2009; Scherrer et al., 2006). In cell culture systems this is frequently accomplished through transfection with fluorescently tagged proteins. However, these *in vitro* systems may not reveal *in vivo* localization. For example, the GPCR melanocortin receptor 4 (MC4R) was recently found to be ciliary only in *in vivo* approaches (Siljee et al., 2018). We sought to use CRISPR/Cas9 mediated genome editing to insert a coding sequence in-frame with a protein, with the understanding that insertion/deletions (in/dels) may be generated in some off-spring, providing an allelic series or additional null alleles from a single round of injections. To test this possibility, we targeted the start codon of *Mchr1,* located in the first of its two exons, to insert the coding sequence for the mCherry fluorescent protein (Shaner et al., 2004). An 836bp PCR product with 57bp homology arms was injected into mouse embryos. The addition of an N-terminal hemagglutinin (HA) signal sequence allows the fusion protein to be threaded through the endoplasmic reticulum, allowing it to maintain its transmembrane spanning properties and properly localize to the membrane (**Figure 1a**) (Lu et al., 2009). From a single round of injections, we obtained a founder animal carrying the mCherry insertion, and 3 other animals with in/dels out of a total of 50 pups screened. The mCherry positive founder was heterozygous for two changes, one allele possessed the correct in-frame insertion, and the other an 8bp deletion creating a presumptive *Mchr1* null allele. The resulting offspring possessed either the mCherry-*Mchr1* fusion allele (hereafter denoted as *Mchr1^ch^*), or the in/del allele as determined by PCR amplification (hereafter denoted as *Mchr1^−^*) (**Figure 1b**). Initial analysis showed that the endogenous mCherry fluorescence could not be detected, and requires antibody amplification (**Figure S1**). Immunostaining with mCherry antibodies revealed co-localization with antibody labeling for MCHR1 in heterozygous and homozygous animals, but not in wildtype littermates (**Figure 1c**). We further find that mCherry labeling co-localizes with adenylate cyclase 3 (ADCY3), a marker for neuronal primary cilia (**Figure S2a**). To confirm ciliary localization, we crossed *Mchr1^+/ch^* mice with mice that possess a conditional loxP flanked intraflagellar transport 88 (*Ift88^F^*) allele, which allows cre mediated cilia ablation (Haycraft et al., 2007). We then targeted cilia loss to GABAergic neurons using a *Gad2^iresCre^* allele (Taniguchi et al., 2011; Ramos et al., 2021). We then assessed ^mCherry^MCHR1 ciliary localization in the nucleus accumbens (NAc), where MCHR1 expression is abundant (Diniz et al., 2020; Ramos et al., 2021; Taniguchi et al., 2011). Heterozygous *Gad2*^iresCre^:*Ift88^+/F^* mice show robust ^mCherry^MCHR1 and ADCY3 cilia co-localization (**Figure S2a**). In homozygous *Gad2*^iresCre^:*Ift88^F/F^* mice, both ciliary ^mCherry^MCHR1 and ADCY3 is largely abolished (**Figure S2b**) (Bowie & Goetz, 2020; Koemeter-Cox et al., 2014; Ramos et al., 2021). These results support the potential utility of the N-terminus tagged MCHR1 allele to assess a specific subcellular localization of a ciliary GPCR *in vivo*.

**Figure 1:**
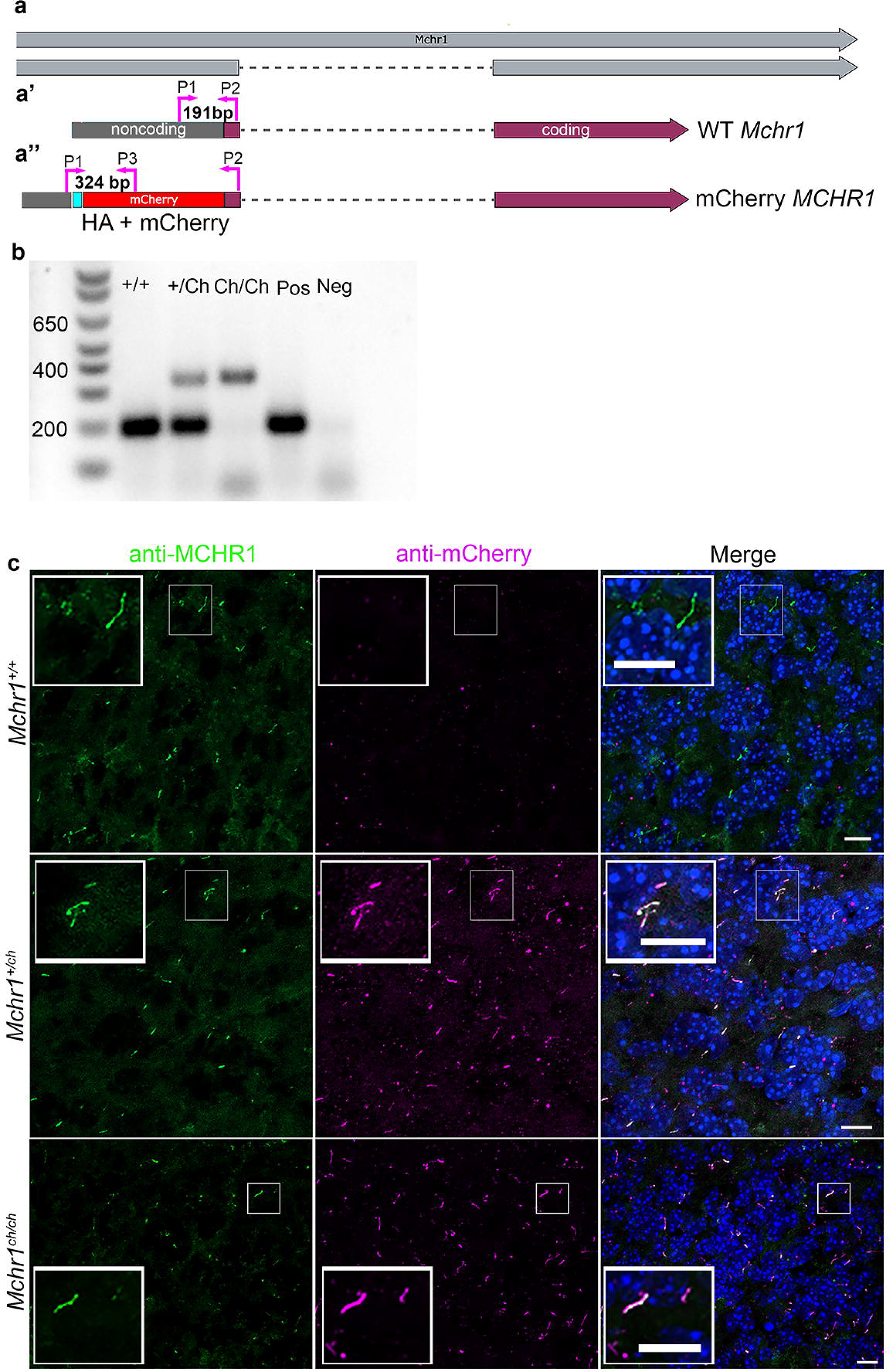
Generation of ^mCherry^Mchr1 N-terminal fusion allele. Targeting *Mchr1* to generate an mCherry fusion allele (**a**) Schematic of genomic targeting with HA tagged mCherry N-terminal start of the coding region of *Mchr1*. Genotyping primers locations are indicated by pink arrows. (**a’)** Mchr1 FOR (P1) and Mchr1 REV (P2) produce a 191bp product from the wildtype (WT) allele, while (**a’’**) Mchr1 FOR (P1) and mCherry REV produce a 324bp product in the modified allele. (**b**) Genomic PCR genotyping for wildtype (*Mchr1^+/+^*), heterozygous (*Mchr1^+/Ch^*), and homozygous (*Mchr1^Ch/Ch^*) mice with C57bl6/j positive control DNA and H_2_O negative control. Molecular weight marker is the first lane on left. (**c**) Immunofluorescence in the olfactory bulb for MCHR1 (green) and mCherry (magenta) in wildtype (*Mchr1^+/+^*), heterozygous (*Mchr1^+/ch^*) and homozygous mCherry (*Mchr1^ch/ch^*). Ciliary co-localization of mCherry and MCHR1 labeling is detected in *Mchr1^+/ch^* and *Mchr1^ch/ch^* mice, but no mCherry labelling is seen in wildtype controls. Nuclei are labeled with DAPI in blue. Scale bars = 10μm

While the lack of robust fluorescence limits the use of ^mCherry^MCHR1 in live imaging approaches, the mCherry epitope provides a tool that allows for labeling with other antibodies and visualization the expression pattern of endogenous levels of this ciliary GPCR. Additionally, by including an HA-signal the protein is properly processed by the cell, and correctly localizes to the cilia. This type of extracellular, N-terminal tag, could therefore improve the ability to perform surface labeling and monitor internalization of ciliary GPCRs following stimulation as well as specific ciliary vs non-ciliary protein-protein interactions through the use of epitope based enrichment approaches like immunoprecipitations (Jenkins et al., 2011; McEwen et al., 2007; Schumacher-Bass et al., 2014). Previous work using viral approaches has found that mCherry tagged ciliary targeting constructs were visible without antibody amplification *in vivo,* suggesting this approach would work (McIntyre et al., 2012; McIntyre, Joiner, Zhang, Iniguez-Lluhi, & Martens, 2015). However, this approach uses the endogenous promotor, and not a viral promoter which may affect expression levels. The need for immunolabeling to visualize ^mCherry^MCHR1 is however consistent with another recent ciliary GPCR fusion mouse model, the MC4R-GFP mouse, in which endogenous fluorescence of GFP could not be visualized, but ciliary labeling was identifiable with GFP antibodies (Siljee et al., 2018). Interestingly, another ciliary GPCR, GPR88, when fused with the brighter fluorescent protein VENUS, can be visualized in cilia without antibody amplification in some regions, although this receptor appears to be ciliary in some cell types and non-ciliary in others (Ehrlich et al., 2018). While ciliary fluorescence is visible in other targeted mouse models with fluorescent C-terminal tags, for example, SSTR3:GFP, Ar13b:GFP, and Arl13b:mCherry, these are all transgenic alleles which may express increased levels of the cilia tagged protein conducive to direct visualization. It is unclear if these types of alleles impinge on normal cilia signaling and functions (Bangs et al., 2015; Delling et al., 2016; O’Connor et al., 2013). Given that ciliary GPCRs may have low levels of endogenous expression, and that mCherry may not be an optimal fluorescent protein, fusion strategies may be more effective with newer, brighter fluorescent proteins (Campbell et al., 2020).

In backcrossing the founder line, allele with the 8bp-deletion in *Mchr1* also resulted in a germline transmission. The small difference in PCR product size can be separated on an agarose gel, however identifying homozygous wildtype and homozygous deletion animals requires product purification followed by sequencing to confirm the deletion (**Figure 2b, c**). Given the small number of base pairs altered, and separation of DNA bands on agarose gels, we have used a breeding strategy of heterozygous (*Mchr1^+/−^*) crossed to homozygous (*Mchr1^−/−^*) mice, as the two bands running together in heterozygous mice produce a visible difference (**Figure 2b)**. To verify that the 8bp deletion was sufficient to induce protein loss, we performed immunostaining with an antibody that recognizes the C-terminal tail of MCHR1. Whereas MCHR1 co-localizes with ADCY3 in *Mchr1^+/−^* mice, no MCHR1 signal is detected in *Mchr1^−/−^* mice (**Figure 2d**). The lack of staining with an antibody to the c-terminal tail provides support that a truncated protein is not produced. In total, the approach to insert a N-terminal fluorescent protein using CRISPR/CAS9 has an added benefit in that it is possible to produce both a reporter line and a knockout mouse line for the same receptor in a single injection attempt. We next sought to determine if N-terminal fusion altered the cilia localization or expression pattern in ^mCherry^MCHR1 mice.

**Figure 2:**
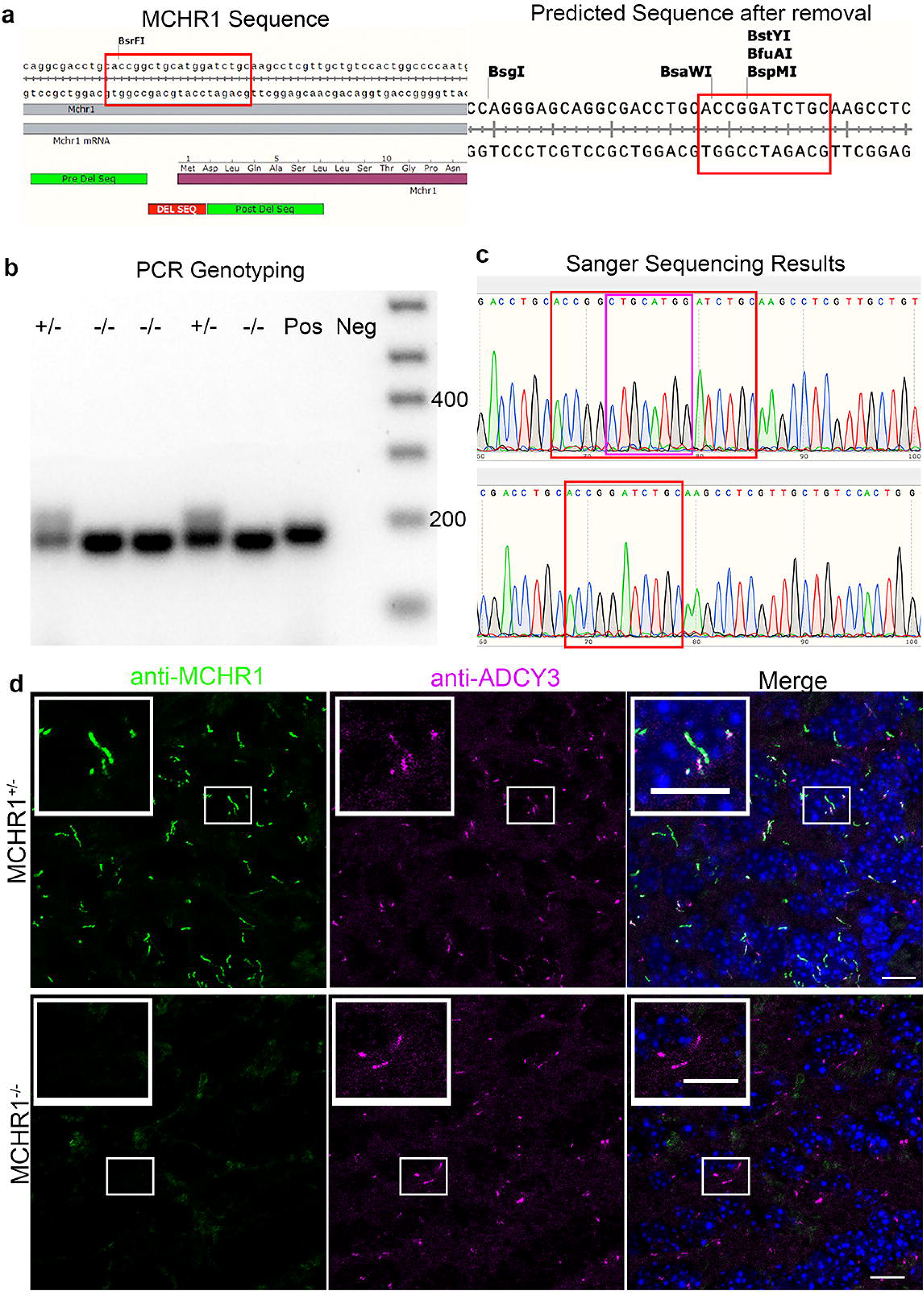
Characterization of an *Mchr1* knockout allele due to an 8 bp deletion. This allele was generated as a by-product of targeting *Mchr1* to insert the coding region for mCherry. (**a**) Schematic of *Mchr1* genomic sequence highlighting the wildtype sequence (left) and the resulting sequence from the deletion (right). Red boxes highlight the reference sequence and the predicted sequence following deletion of the 8bp. (**b**) Genomic PCR gel electrophoresis genotyping, using the *Mchr1* FOR and *Mchr1* REV primers, of animals from *Mchr1^+/−^* x *Mchr1^−/−^* breeding. C57bl6/j positive control DNA (Pos) and H_2_O negative control (Neg) were used. (**c**) Sanger sequence trace files from a *Mchr1^+/+^* mouse (top) and a *Mchr1^−/−^* mouse (bottom) with the red box highlighting the predicted sequences in (a) and the pink box highlighting the 8 deleted base pairs. (**d**) Immunofluorescence for MCHR1 (green) and neuronal cilia protein ADCY3 (magenta) shows loss of staining in *Mchr1^−/−^* brain sections of olfactory bulb compared to *Mchr1^+/−^*. Nuclei are labeled with DAPI in blue. Scale bars = 10μm

### ^mCherry^MCHR1 expression in the brain at different ages

To determine if the *Mchr1^+/ch^* allele maintains an endogenous expression pattern, we performed a temporal analysis in hypothalamic feeding centers of the brain where it is known to be expressed (Engle et al., 2018). ^mCherry^MCHR1 is clearly detected in both the arcuate nucleus (ARC) and paraventricular nucleus (PVN) of the hypothalamus in mice at postnatal day 0 (P0) (**Figure 3a** and **3b**), consistent with MCHR1 localization to neuronal cilia in the hypothalamus (Bansal et al., 2019; Berbari, Johnson, et al., 2008; Berbari, Lewis, Bishop, Askwith, & Mykytyn, 2008; Green, Gu, & Mykytyn, 2012). To assess the localization of ^mcherry^MCHR1 during development and maturation of feeding behaviors, we also imaged brain sections from postnatal day 7 (P7), day 30 (P30), and adult mice (**Figure 3a** and **3b**). ^mCherry^MCHR1 localization is detected in neuronal cilia of both the ARC and PVN neurons at all ages as has been previously reported (Diniz et al., 2020). To confirm the reporter allele was also consistently observed in cilia that express endogenous MCHR1, we co-labeled ARC and PVN sections from *Mchr1^+/ch^* mice with an MCHR1 antibody. At all timepoints in both the ARC and PVN, ^mCherry^MCHR1 efficiently co-labeled the same cilia as the MCHR1 antibody (**Figure 3**) and MCHR1 was not detected in cilia without mCherry colocalization. As immunostaining was done in heterozygous animals, these data confirm complete matching allelic expression. Given the detectable expression from P0 to adult ages, the reporter thus provides a platform for monitoring developmental changes in these neuronal cilia over time as well as under specific behavioral, genetic and pharmacological conditions associated with the MCH signaling axis such as sleep, olfaction, parenting behaviors, and reward (Adams et al., 2011; Ahnaou et al., 2008; Alhassen et al., 2019; Chung et al., 2009; Diniz & Bittencourt, 2017; Hopf, Seif, Chung, & Civelli, 2013; Verret et al., 2003).

**Figure 3:**
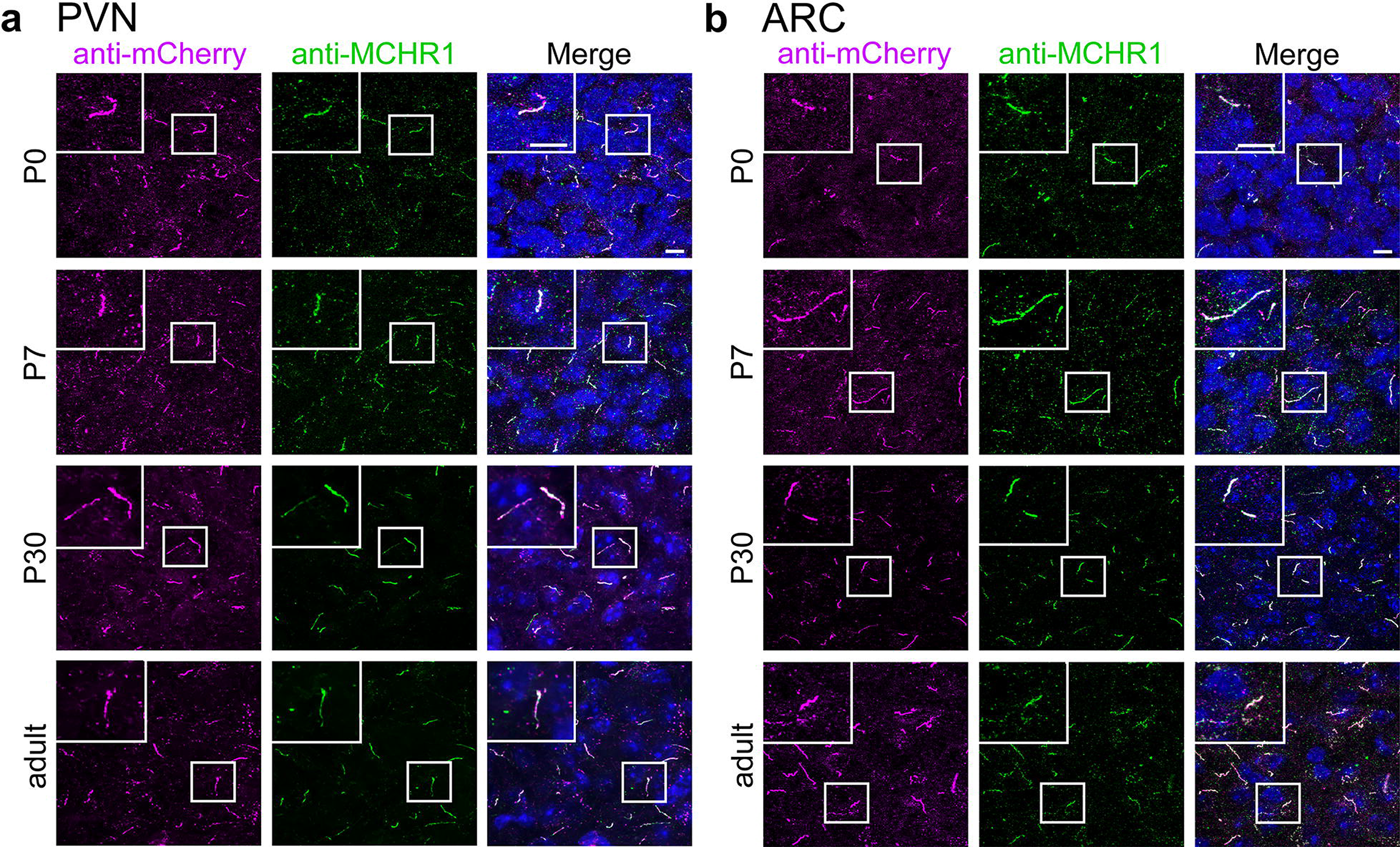
^mCherry^MCHR1 ciliary localization in the developing and adult mouse hypothalamus. Immunofluorescence for mCherry (magenta) and MCHR1 (green) in male *Mchr1^+/ch^* heterozygous hypothalamic section at P0, P7, P30 and Adult (**a**) arcuate nucleus (ARC) and (**b**) paraventricular nucleus (PVN). Hoechst nuclei are blue. Scale bars = 10μm.

Since we saw no overt differences with regards to ^mCherry^MCHR1 localization across development in the hypothalamus, we next focused on assessing localization in other adult brain regions known to express MCHR1. To further examine ^mCherry^MCHR1 expression, we analyzed mCherry localization in the CA1 of the hippocampus, nucleus accumbens (NAc), ventral tegmental area (VTA), olfactory tubercle (OT), olfactory bulb (OB), and in the epithelial cells lining the 3rd ventricle (**Figure 4a-g**). In all of these regions mCherry antibodies labeled cilia projecting from NeuN labeled neuronal cell bodies. In the NAc, ciliary ^mCherry^MCHR1 is highly abundant in neurons. However, in the ventral tegmental area (VTA), co-labeling for the dopaminergic neuron marker tyrosine hydroxylase (TH) reveals sparse labeling of ^mCherry^MCHR1. Further confirming the fidelity of the reporter allele to endogenous expression, mCherry labeling colocalizes with MCHR1 antibody labeling in all regions imaged (**Figure S3**). Taken together these data support the conclusion that ^mCherry^MCHR1 recapitulates the endogenous expression pattern. These results also support a conclusion that MCHR1 is a ciliary enriched GPCR throughout the central nervous system, in contrast to another neuronal ciliary GPCR, GPR88 (Ehrlich et al., 2018). Thus, this new mcherry-*Mchr1* fusion allele can serve as a way to monitor and further identify MCHR1 expression and subcellular localization, particularly when antibody labeling of the endogenous protein is a challenge.

**Figure 4:**
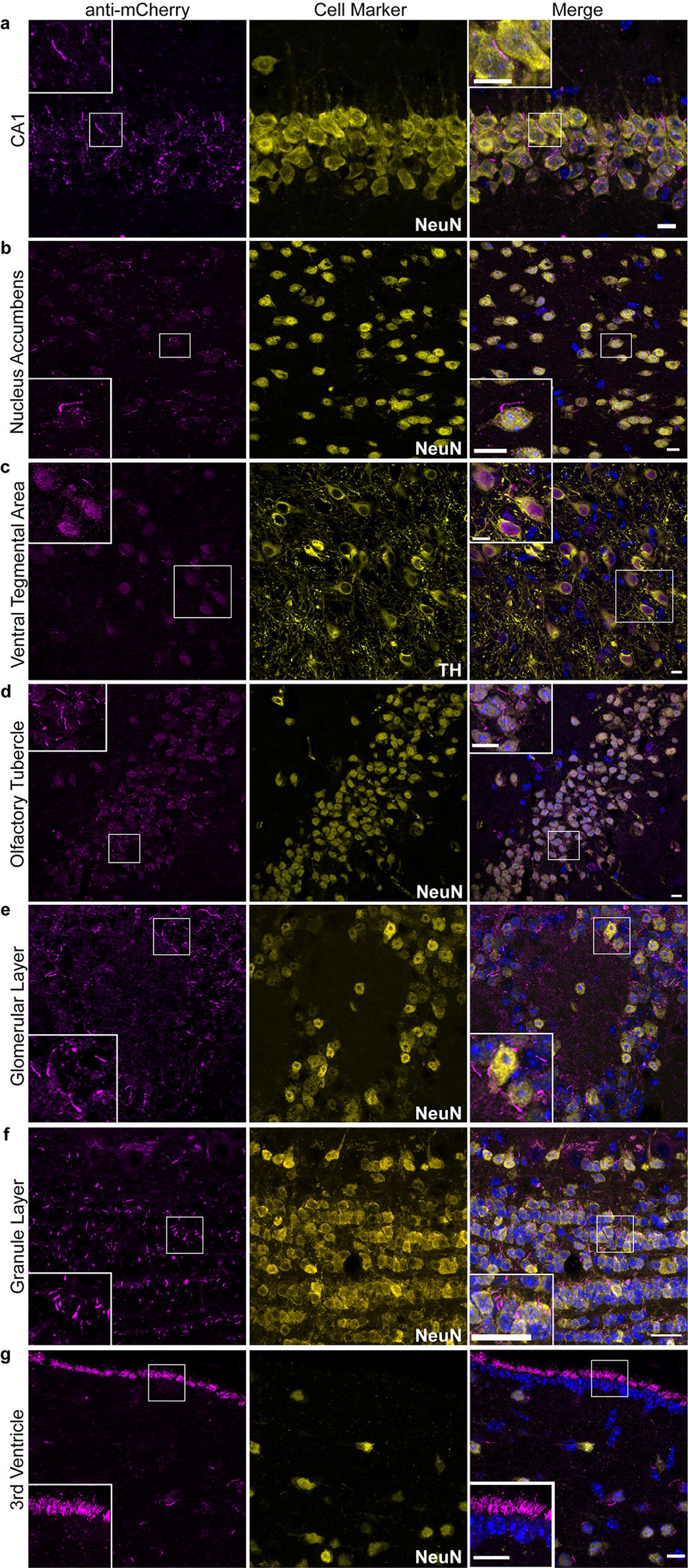
^mCherry^MCHR1 expression matches ciliary localization throughout the adult mouse brain. Immunofluorescence for mCherry (magenta) and neuronal markers NeuN or TH (yellow) in various regions of adult male *Mchr1^+/ch^* sections demonstrate fusion allele cilia localization in (**a**) CA1 region of the hippocampus, (**b**) the nucleus accumbens, (**c**) ventral tegmental area, (**d**) olfactory tubercule, (**e**) glomerular layer of the olfactory bulb (**f**) granule layer of the olfactory bulb and (**g**) motile cilia of the 3^rd^ ventricle. Nuclei are labeled with DAPI in blue. Scale bars = 10μm

### Functional Impact of N-terminal tagged ^mCherry^MCHR1

We then sought to assess potential functional impacts of ^mcherry^MCHR1 on neuronal activity. We performed comparative slice physiology experiments with tissue micro-electrode arrays focused on assessing field recordings from CA1 of the ventral hippocampus where MCHR1 is expressed. We first determined the input/output curves of CA1 following stimulation of CA3 Schaffer collaterals in *Mchr1^+/−^*, *Mchr1^−/−^*, *Mchr1^+/ch^*, and *Mchr1^ch/ch^* mice to assess basal synaptic transmission. The input-output curves demonstrated a significant reduction in activity in *Mchr1^−/−^* mice compared to *Mchr1^+/−^* mice (Two-way ANOVA, main effects of genotype; F(1, 20)=48.77, p <0.0001, input; F(9, 180) = 155.2, p<0.0001, and interaction; F (9, 180) = 34.97, p < 0.0001. n = 5 *Mchr1^+/−^*, *Mchr1^−/−^* mice, 2 slices from each animal) (**Figure 5a**). We also detect a small, but statistically significant reduction in synaptic strength in *Mchr1^ch/ch^* compared to *Mchr1^+/ch^* mice (Two-way ANOVA, main effect of genotype; F(1, 19)=5.541, p <0.0295, input; F(9, 171)=82.74 p<0.0001, and an interaction; F (9, 171) = 2.046, p < 0.0371. n = 5 *Mchr1^+/ch^*, 6 *Mchr1^ch/ch^* mice, 1-2 slices from each animal) (**Figure 5b**). We then tested presynaptic function by determining if paired pulse facilitation (PPF) was altered in *Mchr1^−/−^ and Mchr1^ch/ch^* mice. PPF is a form of short-term synaptic plasticity providing an indirect measure of glutamate release probability at these synapses and changes in PPF have been observed in conditional cilia loss mouse models (Berbari et al., 2014; Koemeter-Cox et al., 2014; Pachoud et al., 2010). The PPF ratio was determined at 4 different inter-stimulus intervals (ISIs: 10, 30, 100, 300ms). The PPF ratio did not differ between *Mchr1^+/−^* and *Mchr1^−/−^* mice across the range of ISI (Two-Way ANOVA, main effect of genotype, F (1, 17) = 0.2434, p = 0.6281, main effect of ISI, F (3,51) = 185.1, p<0.0001. n=5 mice per genotype, 1-2 slices per mouse). (**Figure 5c**). Similarly, the PPF ratio did not differ between *Mchr1^+/ch^* and *Mchr1^ch/ch^* mice (Two-Way ANOVA, main effect of genotype, F(1, 20) = 0.4206, p= 0.524, main effect of ISI, F (3,60) = 245.3, p<0.0001. n=5-6 mice per genotype, 2 slices per mouse) (**Figure 5d**).

**Figure 5:**
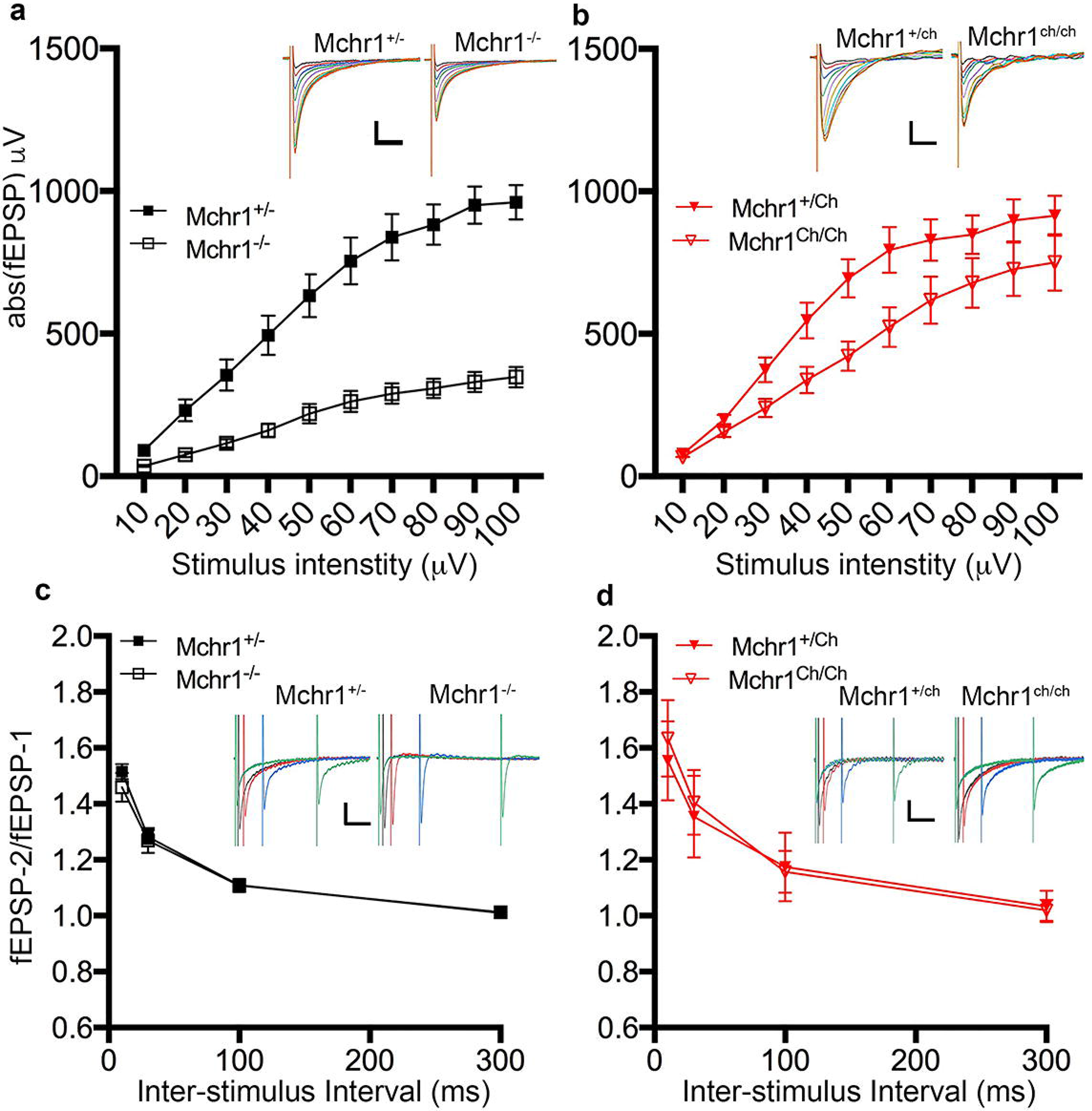
Functional impact on neuronal activity of *Mcrh1^−/−^* and *Mcrh1^ch/ch^* alleles. Electrophysiology was used to evaluate the functional consequence of the 8bp deletion (*Mchr1^−/−^*) and homozygous expression of ^mCherry^MCHR1 (*Mchr1^ch/ch^*). (**a**) Average input/output (I/O) curves from *Mchr^+/−^* (solid squares) and *Mchr1^−/−^* (open squares) mice. Significant differences are detected by Two-way ANOVA with main effects of genotype, F(1, 20)=48.77, p <0.0001, and input strength, F(9, 180) = 155.2, p<0.0001. A significant effect is also detected. F (9, 180) = 34.97, p < 0.0001. n = 5 *Mchr1^+/−^*, *Mchr1^−/−^* mice, 2 slices from each animal). Insets are representative traces from input/output determination using local field potential (LFP) recordings. Scale bar = 200μV/20ms. (**b**) Average input/output (I/O) curves from *Mchr1^+/ch^* (solid triangles) and *Mchr1^ch/ch^* (open triangles) mice. Significant differences are detected by Two-way ANOVA with main effects of genotype, F(1, 19)=5.541, p <0.0295 and input, F(9, 171)=82.74 p<0.0001. An interaction effect is also detected, F (9, 171) = 2.046, p < 0.0371. n = 5 *Mchr1^+/ch^*, 6 *Mchr1^ch/ch^* mice, 1-2 slices from each animal. (**c**) Paired-pulse facilitation (PPF) computed at all inter-stimulus intervals was not significantly different between *Mchr1^+/−^* and *Mchr1^−/−^* mice (Two-Way ANOVA, main effect of genotype, F (1, 17) = 0.2434, p = 0.6281, main effect of ISI, F (3,51) = 185.1, p<0.0001. n=5 mice per genotype, 1-2 slices per mouse). Insets show representative trials. Scale bars = 100μv/100ms. (**d**) Paired-pulse facilitation (PPF) computed at all inter-stimulus intervals was not significantly different between *Mchr1^+/ch^* and *Mchr1^ch/ch^* mice (Two-Way ANOVA, main effect of genotype, F(1, 20) = 0.4206, p= 0.524, main effect of ISI, F (3,60) = 245.3, p<0.0001. n=5-6 mice per genotype, 2 slices per mouse).

The reduction in output strength in *Mchr1^−/−^* mice compared to *Mchr1^+/−^* littermates, with no difference in PPF, is consistent with previously published results using MCHR1 knockout mice (Pachoud et al., 2010). The slight reduction in synaptic strength in *Mchr1^ch/ch^* mice suggests that the fusion of mCherry to the N-terminus reduces some, but not all of the functional activity of MCHR1 in these mice. Whether this is due to a reduction in the amount of the receptor expressed or its functional activity needs to be further determined. Behavioral studies may help elucidate the function of ^mCherry^MCHR1. While MCHR1 knockout mice have normal body weights on regular chow, they were originally found to be resistant to diet induced obesity (Chen et al., 2002; Marsh et al., 2002). We do not detect differences in body weight in either *Mchr1^ch/ch^* or *Mchr1^−/−^* mice, suggesting a similarity there. Future studies looking at diet induced obesity would help clarify the functionality of the fusion ^mCherry^MCHR1 receptor. Together these results show that using an N-terminal fusion approach is effective for monitoring GPCR trafficking and doesn’t alter ciliary localization. Further, in the heterozygous state the expression of ^mCherry^MCHR1 exhibits similar electrophysiological profiles as heterozygous MCHR1 knockout mice suggesting these animals may be suitable to functional studies in which receptor localization can also be studied. The method employed in this paper provided the added benefit of generating a new *Mchr1* knockout allele. Finally, these results do provide a note of caution when either assessing a GPCR tagged allele or an allele used to visualize a cell compartment like the cilium.

## Methods

### Targeting Construct and Mouse Allele Generation

To generate the hemagglutnin (HA)-Signal sequence-mCherry-*Mchr1* mouse we first PCR-amplified an 843bp fragment from the plasmid pENTR/D-topo-HA-mCherry-B2AR. The primer pair - HDR Forward: 5’-ctccactccagggagcaggcgacctgcaccggctgcggccgcatgaagacgatcatc-3’ HDR Reverse: 5’-ggggccagtggacagcaacgaggcttgcagatccataccggtcttgtacagctcgtc-3’ were designed to each consist of 36bp adjacent to the translation initiation codon of *Mchr1* followed by 21 base-pairs of homology in the template mCherry plasmid (Paix et al., 2017). The resulting PCR product with the reporter flanked by 36 base-pair *Mchr1*-homology regions was purified with a Quagen MinElute PCR Purification Kit (28004) for injection. Mouse embryos (from Bl6/SJL F1 hybrid parents), pronuclear injection final concentration: 30ng/ul Cas9 (PNABio)+0.6uM each of crRNA/TracrRNA (guide sequence gctgcatggatctgcaagcc – Dharmacon, Lafayette, CO.). PCR template 5ng/ul. Potential transgenic animals were screened by PCR-amplification using genomic primers outside of the targeting fragment sequence and the products directly sequenced to identify correct homologous insertions of the HA-signal sequence-mCherry reporter at the endogenous *Mchr1* locus (Figure 1a).

### Mice

All procedures were approved by the Institutional Animal Care and Use Committees at the University of Florida and Indiana University-Purdue University Indianapolis. Mice were housed on a standard 12-hour light dark cycle and given food and water *ad libitum*. Mice were weaned and housed with same-sex littermates after postnatal day 21. Ear punches were taken for genotype analysis by polymerase chain reaction.

*Mchr1^+/ch^* founders were crossed to C57Bl6/j mice from the University of Florida colony to generate the colony. Only *Mchr1^+/ch^* mice were used for IHC expression analysis. Experiments used both male and female mice and no differences between sexes were noted. To examine ^mCherry^MCHR1 localization in a cilia knockout model, *Mchr1^+/ch^* mice were bred to a previously generated line crossing *Gad2^iresCre^* (*Gad2^tm2(cre)Zjh^*; Jax stock 010802) mice with floxed *Ift88* (*Ift88^tm1Bky/J^*; Jax stock 022409) mice (Haycraft et al., 2007; Ramos et al., 2021; Taniguchi et al., 2011)

*Mchr1^+/−^* mice were also backcrossed to C57BL6/J (Jax stock 000664) mice. Breeders for experimental animals were *Mchr1^+/−^* mice crossed with *Mchr1^−/−^*. In both lines, homozygous *Mchr1^ch/ch^* and *Mchr1^−/−^* mice are fertile and develop normally. For electrophysiology experiments wildtype mice are *Mchr1^+/−^*. Both male and female mice were used for electrophysiology recordings, but sex differences were not specifically tested.

All Mice were genotyped by extracting DNA from tail clippings with Extracta DNA Prep for PCR – Tissue (Quanta Biosciences) and specified products amplified using 2x KAPA buffer. To genotype *Mch1^+/ch^* mice the following primers were used: *Mchr1* FOR, 5’-gctcaagctccggacaaggc-3’, *Mchr1* REV 5’-caatgtgaaattatcctggccatc-3’, and mCherry REV 5’-ttggtcaccttcagcttgg-3’. With this reaction, the predominate PCR products are 191bp for the wildtype allele (Mchr1 Forward & Mchr1 Reverse) and 324bp for the mCherry modified allele (Mchr1 Forward & mCherry Reverse). However, a 952bp PCR product can be produced by the *Mchr1* primers encompassing the entire mCherry protein. For genotyping *Mchr1^−/−^* knockout mice only *Mchr1* FOR and *Mchr1* REV primers were used. The bands observed for genotypes are a 191bp band for wildtype alleles and 183bp for knockout alleles. To discriminate homozygous wildtype (*Mchr1^+/+^)* from homozygous knockout (*Mchr1^−/−^*), Sanger sequencing using genomic DNA obtained from the tail clippings was used. Genomic DNA was amplified using a Expand High Fidelity PCR protocol (Roche, 11732650001). To prepare for sequencing amplified DNA was gel purified using QIAquick Gel Extraction Kit (QIAGEN, 28706) and prepared according to specifications by GeneWiz (South Plainfield NJ). Sequencing results were analyzed using SnapGene (GSL Biotech, San Diego, CA).

Both strains will be made available to the research community for use.

### Fixation and Tissue Processing for Immunofluorescence

Samples were harvested when mice were P0, P7, P30 and 8-12 weeks old. Mice were anesthetized with 0.1 ml/ 10 g of body weight dose of 2.0% tribromoethanol (Sigma Aldrich, St. Louis, MO) and transcardially perfused with PBS followed by 4% paraformaldehyde (Affymetrix Inc., Cleveland, OH). Brains were postfixed in 4% paraformaldehyde for 24 hours at 4°C and then cryoprotected using sucrose in PBS for 16–24 hours. Cryoprotected brains were embedded in Optimal Cutting Temperature compound (Fisher Healthcare, Houston, TX) and sectioned in a freezing cryostat at a thickness of 10-35 μm.

### Immunofluorescence

Immunofluorescence experiments were carried out in both labs using differing methods for detection of primary cilia was performed as previously described (Bansal et al., 2019). Briefly, blocking was done with PBS containing either 2% Donkey or Goat serum, 0.3% titron X-100, 1% BSA, and 0.02% Sodium azide (Dot Scientific) for 30-60 minutes in a moist chamber. After incubation, primary antibodies diluted in blocking solution were added to sections and slides were cover slipped using parafilm and incubated overnight at 4°C in a humidified chamber. On day 2, after three 5-minutes washes, secondary antibodies diluted in blocking were added to sections and slides were incubated for 60 min away from light. After two washes, nuclei were stained with Hoechst 33342 1:1000 (Thermo Fisher Scientific, Waltham, MA, United States) and slides were cover slipped using slow fade prolong antifade (Invitrogen, Carlsbad, CA, United States) and sealed with nail polish. Primary cilia were stained using anti-adenylate cyclase III (ADCY3) (mouse, EnCor MCA-1A12 1:1000; rabbit, EnCor RPCA-ACIII 1:2000), anti-MCHR1 (rabbit, Invitrogen 711649 1:250) and anti-mCherry (chicken, NOVUS NBP2-25158 1:250; chicken, EnCor CPCA-mCherry 1:1000) antibodies. Cells were stained using anti-NeuN (mouse, Millipore MAB377 1:1000) and anti-tyrosine hydroxylase (mouse, R&D Systems MAB7566 1:500). Fluorescent-conjugated secondary antibodies (ThermoFisher Scientific, Catalog numbers, goat anti-rabbit 488 A-27034; goat anti-rabbit 594, A-11012; goat anti-mouse IgG1 594, A-21225; goat anti-mouse IgG2a 488, A-21131; goat anti-mouse IgG2a 594, A-21135; goat anti-chicken 488, A-11039) were applied (at 1:1000 dilution). Additional secondary antibodies include Alexa Fluor 488-conjugated AffinPure Donkey Anti-Chicken (1:800)(Jackson ImmunResearch 703-545-155), and Alexa Fluor 647 donkey anti-rabbit (1:1000)(Thermo Fisher Scientific A31573).

### Imaging

Images were captured using a Leica SP8 confocal microscope in resonant scanning mode in the Berbari lab, or with a Nikon TiE-PFS-A1R confocal microscope equipped with a 488 nm laser diode with a 510-560 nm band pass filter, and a 561nm laser with a 575-625 nm band pass filter in the McIntyre lab. Confocal *Z*-stacks were processed with Nikon Elements software and NIH ImageJ software.

### Slice Preparation and Extracellular recordings

For extracellular recordings adult (4-7 weeks) male and female *Mchr1^+/−^*, *Mchr1^−/−^*, *Mchr1^+/ch^*, and *Mchr1^ch/ch^* mice were deeply anesthetized with isofluorane and then decapitated. Brains were removed into ice◻cold cutting artificial cerebrospinal fluid (ACSF) containing 124 mM NaCl, 26 mM NaHCO_3_, 3 mM KCl, 1.25 mM KH_2_PO_4_, 2 mM CaCl_2_, 10 mM glucose, and 1 mM MgSO_4_ bubbled with 95% O_2_/5% CO_2_. Coronal hippocampal slices were made (300 μM) using a Leica VT 1000S (Leica Biosystems, Wetzlar, Germany) vibrating microtome. Slices were incubated at 37°C in recording ACSF (124 mM NaCl, 26 mM NaHCO_3_, 3 mM KCl, 1.25 mM KH_2_PO_4_, 2 mM CaCl_2_, 10 mM glucose, and 1 mM MgSO_4_) bubbled with 95% O_2_/5% CO_2_ for 1 hr prior to recording.

Slices were placed onto MED‐P501A 16‐channel multielectrode arrays (Alpha MED Scientific Inc., Osaka JP) with the CA1 pyramidal cell layer positioned directly over the array contacts. Recordings were made at 32°C in recirculating recording ACSF bubbled with 95% O_2_/5% CO_2_. Evoked potentials were captured using MED64 hardware and Mobius software (MED64 system, Alpha MED Scientific Inc., Osaka JP). Synaptic field potentials were evoked by stimulation of the SC pathway at a contact located in stratum radiatum. For every slice, we analyzed data from a recording channel in CA1 that was representative of responses across multiple channels within the same slice as to avoid biases in choosing responses at the extremes. Using data from the input/output (IO) curve, stimulus intensity was determined for paired pulse facilitation (PPF) experiments as the input that produced an evoked field response amplitude 30-40% of the maximum response (generally 20-40 μA, 0.2 ms). IO curves were generated by measuring field excitatory postsynaptic potentials (fEPSPs) following stimulus inputs ranging from −10 to −100 μV in 10 μV increments. PPF ratios were measured for 10, 30, 100 and 300 ms inter-stimulus intervals and determined by dividing the second fEPSP (fEPSP-2) by the first fEPSP (fEPSP-1).

## Acknowledgments

JCM, HK, and RRR designed and produced the MCHR1 targeting constructs. KRJ, TK, RB, SE performed IHC experiments. AZ and TE performed extracellular electrophysiological recordings. SE, NFB, and JCM analyzed data and wrote the manuscript with edits from all authors. The authors would like to thank the UF Cell and Tissue Analysis Core for confocal microscopy assistance (NIH grant 1S10OD020026), Johns Hopkins University facility for assistance in producing founder mice and the Berbari and McIntyre Labs for critical reading of the manuscript.

## Supplemental Figure Legend

**Supplemental Figure 1: Endogenous fluorescence ^mCherry^MCHR1 is not detected** Immunofluorescence image of both antibody-labeled and unlabeled mCherry. Whereas the mCherry antibody is able to bind to the protein as detected in the 488 channel, endogenous fluorescence is not detected in the 547 channel. Labeling for MCHR1 specifically reveals co-localization with antibody labeled mCherry. Scale bar = 10μm Hoechst nuclei are blue.

**Supplemental Figure 2: IFT88 loss disrupts ^mCherry^MCHR1 ciliary localization.** Immunofluorescence staining for mCherry and ADCY3 in the nucleus accumbens, where endogenous MCHR1 expression is high, of *Mchr1^+/ch^:Gad2^iresCre^:If88^+/F^* and *Mchr1+/ch:Gad2^iresCre^:Ift88^F/F^* mice. (**a**) In wildtype mice (*If88^+/F^*), ^mCherry^MCHR1 and ADCY3 are co-localized in cilia structures. (**b**) In IFT88 knockout mice (*Ift88^F/F^*), ciliary labeling of both mCherry and ADCY3 is absent in neurons expressing *Gad2^iresCre^*. DAPI nuclei are blue. Scale bars = 10μm

**Supplemental Figure 3: ^mCherry^MCHR1 expression matches MCHR1 localization throughout the adult mouse brain.** Immunofluorescence for mCherry (magenta) and MCHR1 (green) in male *Mchr1^+/ch^* adult mice in (**a**) CA1 of hippocampus (CA1) and (**b**) nucleus accumbens. Hoechst nuclei are blue. Scale bars = 10μm.

## References

Adams, A. C., Domouzoglou, E. M., Chee, M. J., Segal-Lieberman, G., Pissios, P., & Maratos-Flier, E. (2011). Ablation of the hypothalamic neuropeptide melanin concentrating hormone is associated with behavioral abnormalities that reflect impaired olfactory integration. Behav Brain Res, 224(1), 195–200. doi:10.1016/j.bbr.2011.05.039

Ahnaou, A., Drinkenburg, W. H., Bouwknecht, J. A., Alcazar, J., Steckler, T., & Dautzenberg, F. M. (2008). Blocking melanin-concentrating hormone MCH1 receptor affects rat sleep-wake architecture. Eur J Pharmacol, 579(1-3), 177–188. doi:10.1016/j.ejphar.2007.10.017

Alhassen, L., Phan, A., Alhassen, W., Nguyen, P., Lo, A., Shaharuddin, H., Alachkar, A. (2019). The role of Olfaction in MCH-regulated spontaneous maternal responses. Brain Res, 1719, 71–76. doi:10.1016/j.brainres.2019.05.021

Bangs, F. K., Schrode, N., Hadjantonakis, A. K., & Anderson, K. V. (2015). Lineage specificity of primary cilia in the mouse embryo. Nat Cell Biol, 17(2), 113–122. doi:10.1038/ncb3091

Bansal, R., Engle, S. E., Antonellis, P. J., Whitehouse, L. S., Baucum, A. J., 2nd, Cummins, T. R., Berbari, N. F. (2019). Hedgehog Pathway Activation Alters Ciliary Signaling in Primary Hypothalamic Cultures. Front Cell Neurosci, 13, 266. doi:10.3389/fncel.2019.00266

Berbari, N. F., Johnson, A. D., Lewis, J. S., Askwith, C. C., & Mykytyn, K. (2008). Identification of ciliary localization sequences within the third intracellular loop of G protein-coupled receptors. Mol Biol Cell, 19(4), 1540–1547. doi:10.1091/mbc.E07-09-0942

Berbari, N. F., Lewis, J. S., Bishop, G. A., Askwith, C. C., & Mykytyn, K. (2008). Bardet-Biedl syndrome proteins are required for the localization of G protein-coupled receptors to primary cilia. Proc Natl Acad Sci U S A, 105(11), 4242–4246. doi:10.1073/pnas.0711027105

Berbari, N. F., Malarkey, E. B., Yazdi, S. M., McNair, A. D., Kippe, J. M., Croyle, M. J., Yoder, B. K. (2014). Hippocampal and cortical primary cilia are required for aversive memory in mice. PLoS One, 9(9), e106576. doi:10.1371/journal.pone.0106576

Berbari, N. F., O’Connor, A. K., Haycraft, C. J., & Yoder, B. K. (2009). The primary cilium as a complex signaling center. Curr Biol, 19(13), R526–535. doi:10.1016/j.cub.2009.05.025

Bittencourt, J. C., Presse, F., Arias, C., Peto, C., Vaughan, J., Nahon, J. L., Sawchenko, P. E. (1992). The melanin-concentrating hormone system of the rat brain: an immuno- and hybridization histochemical characterization. J Comp Neurol, 319(2), 218–245. doi:10.1002/cne.903190204

Bowie, E., & Goetz, S. C. (2020). TTBK2 and primary cilia are essential for the connectivity and survival of cerebellar Purkinje neurons. Elife, 9. doi:10.7554/eLife.51166

Campbell, B. C., Nabel, E. M., Murdock, M. H., Lao-Peregrin, C., Tsoulfas, P., Blackmore, M. G., Petsko, G. A. (2020). mGreenLantern: a bright monomeric fluorescent protein with rapid expression and cell filling properties for neuronal imaging. Proc Natl Acad Sci U S A, 117(48), 30710–30721. doi:10.1073/pnas.2000942117

Chan, F., Bradley, A., Wensel, T. G., & Wilson, J. H. (2004). Knock-in human rhodopsin-GFP fusions as mouse models for human disease and targets for gene therapy. Proc Natl Acad Sci U S A, 101(24), 9109–9114. doi:10.1073/pnas.0403149101

Chen, Y., Hu, C., Hsu, C. K., Zhang, Q., Bi, C., Asnicar, M., Shi, Y. (2002). Targeted disruption of the melanin-concentrating hormone receptor-1 results in hyperphagia and resistance to diet-induced obesity. Endocrinology, 143(7), 2469–2477. doi:10.1210/endo.143.7.8903

Chung, S., Hopf, F. W., Nagasaki, H., Li, C. Y., Belluzzi, J. D., Bonci, A., & Civelli, O. (2009). The melanin-concentrating hormone system modulates cocaine reward. Proc Natl Acad Sci U S A, 106(16), 6772–6777. doi:10.1073/pnas.0811331106

Delling, M., Indzhykulian, A. A., Liu, X., Li, Y., Xie, T., Corey, D. P., & Clapham, D. E. (2016). Primary cilia are not calcium-responsive mechanosensors. Nature, 531(7596), 656–660. doi:10.1038/nature17426

Diniz, G. B., Battagello, D. S., Klein, M. O., Bono, B. S. M., Ferreira, J. G. P., Motta-Teixeira, L. C., Bittencourt, J. C. (2020). Ciliary melanin-concentrating hormone receptor 1 (MCHR1) is widely distributed in the murine CNS in a sex-independent manner. J Neurosci Res, 98(10), 2045–2071. doi:10.1002/jnr.24651

Diniz, G. B., & Bittencourt, J. C. (2017). The Melanin-Concentrating Hormone as an Integrative Peptide Driving Motivated Behaviors. Front Syst Neurosci, 11, 32. doi:10.3389/fnsys.2017.00032

Drake, M. T., Shenoy, S. K., & Lefkowitz, R. J. (2006). Trafficking of G protein-coupled receptors. Circ Res, 99(6), 570–582. doi:10.1161/01.RES.0000242563.47507.ce

Ehrlich, A. T., Semache, M., Bailly, J., Wojcik, S., Arefin, T. M., Colley, C., Kieffer, B. L. (2018). Mapping GPR88-Venus illuminates a novel role for GPR88 in sensory processing. Brain Struct Funct, 223(3), 1275–1296. doi:10.1007/s00429-017-1547-3

Engle, S. E., Antonellis, P. J., Whitehouse, L. S., Bansal, R., Emond, M. R., Jontes, J. D., Berbari, N. F. (2018). A CreER mouse to study melanin concentrating hormone signaling in the developing brain. Genesis, 56(8), e23217. doi:10.1002/dvg.23217

Green, J. A., Gu, C., & Mykytyn, K. (2012). Heteromerization of ciliary G protein-coupled receptors in the mouse brain. PLoS One, 7(9), e46304. doi:10.1371/journal.pone.0046304

Green, J. A., Schmid, C. L., Bley, E., Monsma, P. C., Brown, A., Bohn, L. M., & Mykytyn, K. (2016). Recruitment of beta-Arrestin into Neuronal Cilia Modulates Somatostatin Receptor Subtype 3 Ciliary Localization. Mol Cell Biol, 36(1), 223–235. doi:10.1128/MCB.00765-15

Haycraft, C. J., Zhang, Q., Song, B., Jackson, W. S., Detloff, P. J., Serra, R., & Yoder, B. K. (2007). Intraflagellar transport is essential for endochondral bone formation. Development, 134(2), 307–316. doi:10.1242/dev.02732

Hopf, F. W., Seif, T., Chung, S., & Civelli, O. (2013). MCH and apomorphine in combination enhance action potential firing of nucleus accumbens shell neurons in vitro. PeerJ, 1, e61. doi:10.7717/peerj.61

Jenkins, P. M., McIntyre, J. C., Zhang, L., Anantharam, A., Vesely, E. D., Arendt, K. L., Martens, J. R. (2011). Subunit-dependent axonal trafficking of distinct alpha heteromeric potassium channel complexes. J Neurosci, 31(37), 13224–13235. doi:10.1523/JNEUROSCI.0976-11.2011

Koemeter-Cox, A. I., Sherwood, T. W., Green, J. A., Steiner, R. A., Berbari, N. F., Yoder, B. K., Mykytyn, K. (2014). Primary cilia enhance kisspeptin receptor signaling on gonadotropin-releasing hormone neurons. Proc Natl Acad Sci U S A, 111(28), 10335–10340. doi:10.1073/pnas.1403286111

Kroeze, W. K., Sheffler, D. J., & Roth, B. L. (2003). G-protein-coupled receptors at a glance. J Cell Sci, 116(Pt 24), 4867–4869. doi:10.1242/jcs.00902

Lu, R., Li, Y., Zhang, Y., Chen, Y., Shields, A. D., Winder, D. G., Wang, Q. (2009). Epitope-tagged receptor knock-in mice reveal that differential desensitization of alpha2-adrenergic responses is because of ligand-selective internalization. J Biol Chem, 284(19), 13233–13243. doi:10.1074/jbc.M807535200

Marinissen, M. J., & Gutkind, J. S. (2001). G-protein-coupled receptors and signaling networks: emerging paradigms. Trends Pharmacol Sci, 22(7), 368–376. doi:10.1016/s0165-6147(00)01678-3

Marsh, D. J., Weingarth, D. T., Novi, D. E., Chen, H. Y., Trumbauer, M. E., Chen, A. S., Qian, S. (2002). Melanin-concentrating hormone 1 receptor-deficient mice are lean, hyperactive, and hyperphagic and have altered metabolism. Proc Natl Acad Sci U S A, 99(5), 3240–3245. doi:10.1073/pnas.052706899

McEwen, D. P., Schumacher, S. M., Li, Q., Benson, M. D., Iniguez-Lluhi, J. A., Van Genderen, K. M., & Martens, J. R. (2007). Rab-GTPase-dependent endocytic recycling of Kv1.5 in atrial myocytes. J Biol Chem, 282(40), 29612–29620. doi:10.1074/jbc.M704402200

McIntyre, J. C., Davis, E. E., Joiner, A., Williams, C. L., Tsai, I. C., Jenkins, P. M., Martens, J. R. (2012). Gene therapy rescues cilia defects and restores olfactory function in a mammalian ciliopathy model. Nat Med, 18(9), 1423–1428. doi:10.1038/nm.2860

McIntyre, J. C., Hege, M. M., & Berbari, N. F. (2016). Trafficking of ciliary G protein-coupled receptors. Methods Cell Biol, 132, 35–54. doi:10.1016/bs.mcb.2015.11.009

McIntyre, J. C., Joiner, A. M., Zhang, L., Iniguez-Lluhi, J., & Martens, J. R. (2015). SUMOylation regulates ciliary localization of olfactory signaling proteins. J Cell Sci, 128(10), 1934–1945. doi:10.1242/jcs.164673

Michel, M. C., Wieland, T., & Tsujimoto, G. (2009). How reliable are G-protein-coupled receptor antibodies? Naunyn Schmiedebergs Arch Pharmacol, 379(4), 385–388. doi:10.1007/s00210-009-0395-y

O’Connor, A. K., Malarkey, E. B., Berbari, N. F., Croyle, M. J., Haycraft, C. J., Bell, P. D., Yoder, B. K. (2013). An inducible CiliaGFP mouse model for in vivo visualization and analysis of cilia in live tissue. Cilia, 2(1), 8. doi:10.1186/2046-2530-2-8

Pachoud, B., Adamantidis, A., Ravassard, P., Luppi, P. H., Grisar, T., Lakaye, B., & Salin, P. A. (2010). Major impairments of glutamatergic transmission and long-term synaptic plasticity in the hippocampus of mice lacking the melanin-concentrating hormone receptor-1. J Neurophysiol, 104(3), 1417–1425. doi:10.1152/jn.01052.2009

Paix, A., Folkmann, A., Goldman, D. H., Kulaga, H., Grzelak, M. J., Rasoloson, D., Seydoux, G. (2017). Precision genome editing using synthesis-dependent repair of Cas9-induced DNA breaks. Proc Natl Acad Sci U S A, 114(50), E10745–E10754. doi:10.1073/pnas.1711979114

Patwardhan, A., Cheng, N., & Trejo, J. (2021). Post-Translational Modifications of G Protein-Coupled Receptors Control Cellular Signaling Dynamics in Space and Time. Pharmacol Rev, 73(1), 120–151. doi:10.1124/pharmrev.120.000082

Ramos, C., Roberts, J. B., Jasso, K. R., Ten Eyck, T. W., Everett, T., Pozo, P., McIntyre, J. C. (2021). Neuron-specific cilia loss differentially alters locomotor responses to amphetamine in mice. J Neurosci Res, 99(3), 827–842. doi:10.1002/jnr.24755

Saito, Y., Cheng, M., Leslie, F. M., & Civelli, O. (2001). Expression of the melanin-concentrating hormone (MCH) receptor mRNA in the rat brain. J Comp Neurol, 435(1), 26–40.

Scherrer, G., Tryoen-Toth, P., Filliol, D., Matifas, A., Laustriat, D., Cao, Y. Q., Kieffer, B. L. (2006). Knockin mice expressing fluorescent delta-opioid receptors uncover G protein-coupled receptor dynamics in vivo. Proc Natl Acad Sci U S A, 103(25), 9691–9696. doi:10.1073/pnas.0603359103

Schou, K. B., Pedersen, L. B., & Christensen, S. T. (2015). Ins and outs of GPCR signaling in primary cilia. EMBO Rep, 16(9), 1099–1113. doi:10.15252/embr.201540530

Schumacher-Bass, S. M., Vesely, E. D., Zhang, L., Ryland, K. E., McEwen, D. P., Chan, P. J., Martens, J. R. (2014). Role for myosin-V motor proteins in the selective delivery of Kv channel isoforms to the membrane surface of cardiac myocytes. Circ Res, 114(6), 982–992. doi:10.1161/CIRCRESAHA.114.302711

Shaner, N. C., Campbell, R. E., Steinbach, P. A., Giepmans, B. N., Palmer, A. E., & Tsien, R. Y. (2004). Improved monomeric red, orange and yellow fluorescent proteins derived from Discosoma sp. red fluorescent protein. Nat Biotechnol, 22(12), 1567–1572. doi:10.1038/nbt1037

Shinde, S. R., Nager, A. R., & Nachury, M. V. (2020). Ubiquitin chains earmark GPCRs for BBSome-mediated removal from cilia. J Cell Biol, 219(12). doi:10.1083/jcb.202003020

Siljee, J. E., Wang, Y., Bernard, A. A., Ersoy, B. A., Zhang, S., Marley, A., Vaisse, C. (2018). Subcellular localization of MC4R with ADCY3 at neuronal primary cilia underlies a common pathway for genetic predisposition to obesity. Nat Genet, 50(2), 180–185. doi:10.1038/s41588-017-0020-9

Taniguchi, H., He, M., Wu, P., Kim, S., Paik, R., Sugino, K., Huang, Z. J. (2011). A resource of Cre driver lines for genetic targeting of GABAergic neurons in cerebral cortex. Neuron, 71(6), 995–1013. doi:10.1016/j.neuron.2011.07.026

Verret, L., Goutagny, R., Fort, P., Cagnon, L., Salvert, D., Leger, L., Luppi, P. H. (2003). A role of melanin-concentrating hormone producing neurons in the central regulation of paradoxical sleep. BMC Neurosci, 4, 19. doi:10.1186/1471-2202-4-19

Ye, F., Nager, A. R., & Nachury, M. V. (2018). BBSome trains remove activated GPCRs from cilia by enabling passage through the transition zone. J Cell Biol, 217(5), 1847–1868. doi:10.1083/jcb.201709041

