## Supplementary figures and images for "An N-terminal Fusion Allele to Study Melanin Concentrating Hormone Receptor 1"

### Supplemental figure 1

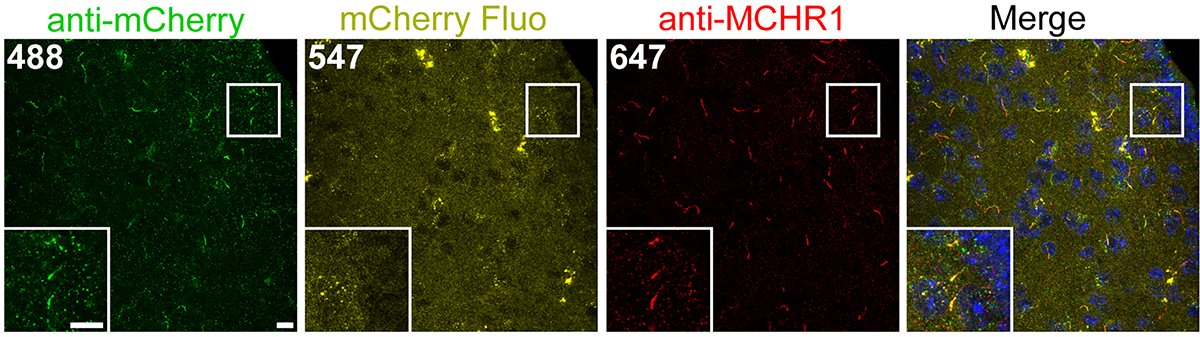

### Supplemental figure 2

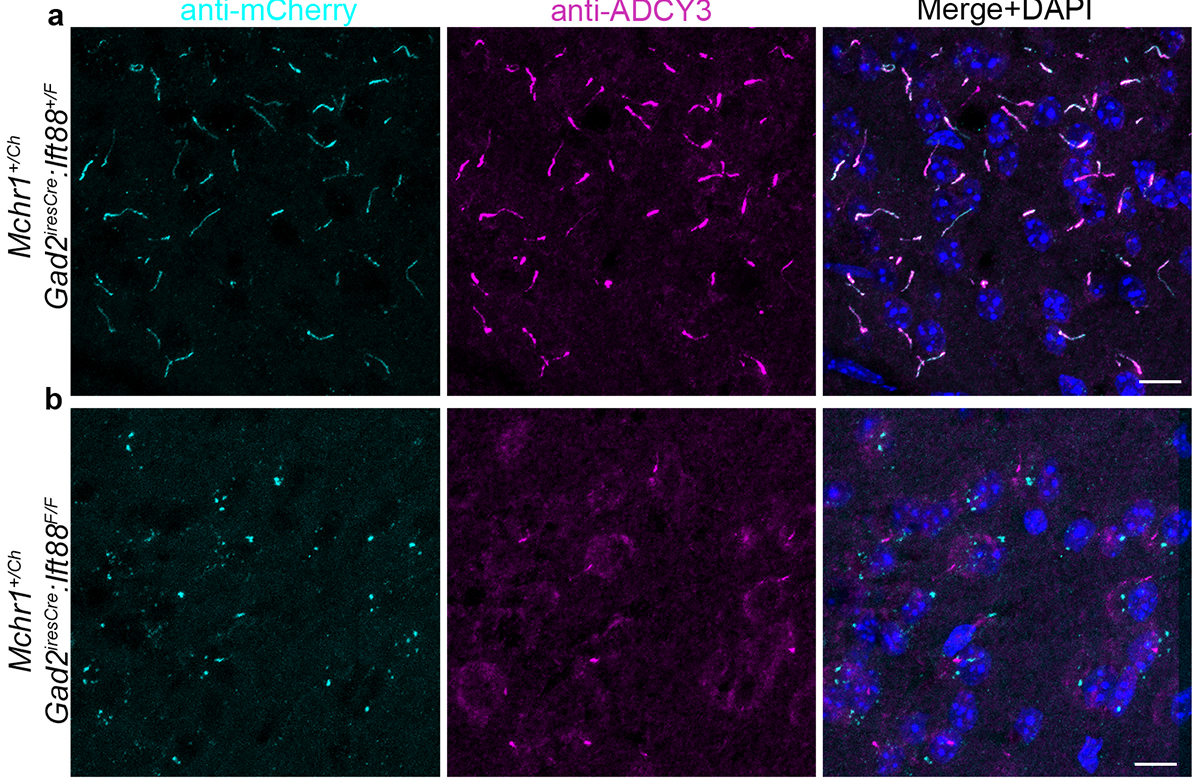

### Supplemental Figure 3

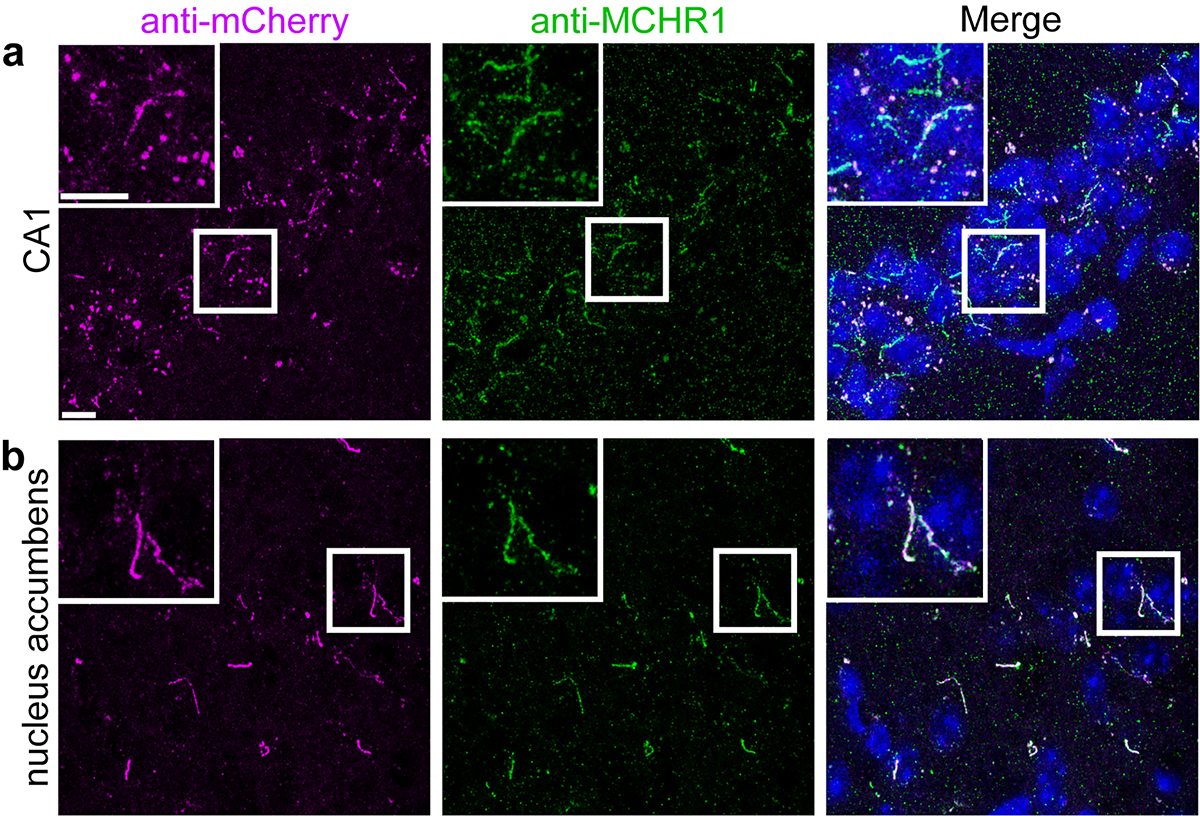
